# Random-forest segmentation and spatial analysis of injected cardiac spheroids in optically cleared myocardium

**DOI:** 10.64898/2026.03.20.713110

**Authors:** Ahmed Elnageh, Sasha Forbes, Steven M. Moreno, Sharika Mohanan, Godfrey Smith, Eline Huethorst, Caroline Müllenbroich

## Abstract

Accurate quantification of transplanted cardiac spheroids requires three-dimensional localisation within intact myocardium, yet this remains technically challenging. Optical clearing and light-sheet microscopy enable volumetric imaging of injection sites, but automated segmentation is difficult when transplanted spheroids and host tissue are labelled with the same fluorescent markers and cannot be separated by simple thresholding. We developed a random forest based pixel classification workflow for 3D detection of injected hiPSC derived cardiomyocyte and H9c2 spheroids in optically cleared rabbit myocardium. A supervised classifier trained on intensity, edge, and texture features generated a segmentation then grouped pixels via connected component analysis to reconstruct individual spheroids. The method showed good agreement with manual annotation and enabled automated extraction of spheroid size and spatial metrics. This accessible workflow enables reproducible three–dimensional quantification of transplanted spheroids in large light-sheet microscopy datasets and provides a practical route from volumetric imaging to spatial metrics in cardiac regeneration studies.

## 1 INTRODUCTION

Following myocardial infarction, cardiac function declines due to the heart’s limited intrinsic regenerative capacity [1, 2]. Regenerative strategies therefore aim to replace dead cardiomyocytes with healthy cardiomyocytes such as human induced pluripotent stem-cell derived cardiomyocytes (hiPSC-CMs) often by epicardial grafting of engineered heart tissues [3, 4, 5] or injection of cell spheroids into the myocardium [6]. Beyond demonstrating graft viability, graft-host coupling and thus therapeutic efficacy depend critically on post-injection localisation, retention, clustering and penetration depth of injected cells. Quantifying the spatial distribution of injected spheroids within intact tissue remains challenging. Conventional histology is destructive, sparsely sampling and unable to resolve full injection tracks. Volumetric imaging approaches are required to characterise spheroid position and organisation within whole tissue sections in order to investigate the graft-host interface. Optical clearing [7] combined with meso-scale light-sheet microscopy [8, 9] enables imaging of large cardiac volumes at cellular resolution [10]. This approach can visualise injected material across millimetre-scales regions with preserved anatomical context, however, automated segmentation is needed to provide quantitative metrics and insight into retention and dispersion of transplanted spheroids. Segmentation is non-trivial because injected spheroids often share fluorescent labels with surrounding myocardium, making intensity-based thresholding unreliable. Structural heterogeneity, depth-dependent signal variation and the presence of bright features such as vasculature further complicate analysis. The resulting large volumetric datasets make manual annotation impractical and subjective. Although deep learning methods [11, 12] can achieve high performance, they require substantial annotated datasets and computational resources, while simpler threshold- or morphology-based approaches lack robustness in heterogeneous cleared tissue. Random forest pixel classification [13, 14] offers a flexible and computationally lightweight alternative. A random forest is a machine learning algorithm consisting of an ensemble of decision trees which classify pixels based on a set of features extracted from the image. By aggregating the predictions of numerous trees, the variance of the predictions is reduced and the predictions can be efficient even with modest training datasets. Importantly the aggregation of multiple trees allows the algorithm to handle non-linear decision boundaries well, which is vital for accurate decisions in highly heterogenous images. Here, we develop and validate a random forest-based workflow for automated detection and 3D reconstruction of injected spheroids in optically cleared myocardial tissue imagined with light-sheet microscopy. By integrating multiscale feature-based pixel classification with object-level reconstruction we extract quantitative spatial metrics including spheroid size, nearest-neighbour distance and depth relative to the epicardium. We demonstrate the approach on hiPSC-derived cardiomyocyte and H9c2 spheroids in cleared rabbit heart tissue, establishing a practical and reproducible pipeline for quantitative analysis in cardiac regenerative studies.

## 2 METHODS

### 2.1 Sample preparation and imaging

Commercial human induced pluripotent stem cell-derived cardiomyocytes (hiPSC–CMs) (Celogics, Cat# C50) or H9c2(2-1) rat myoblasts (ATCC, Cat#CRL01446) were used to generate spheroids which formed by self-aggregation (mean diameter of 340 µm) after seeding in Aggrewell 800 microwell plates. Day 3-5 after plating, spheroids were injected into the left ventricle of a Langendorff-perfused rabbit heart (New Zealand White), in accordance with the UK Animals Scientific Procedures Act 1986 and under the Project License (PP5254544) for a study on cell therapy for cardiac regeneration. Each injection site was marked with a suture and after functional studies, the injection region was cut, fixed in 4% paraformaldehyde (PFA) for 5-7 days at 4 ^◦^C, before being washed in PBS and cleared according to the CUBIC Stars protocol [15]. Samples were placed in 50% CUBIC-L (with dH20) for 24 hours, then placed in 100% CUBIC-L for 4 weeks. After clearing, all samples were stained with the cell membrane stain Wheat Germ Agglutin (WGA) Alexa Fluor 647 (1:100, 10µg/ml) (Invitrogen, Cat#W32466) and the nuclear stain SYTOX Green (1:2500, 2µM) (ThermoFisher, Cat#S7020) in PBST (0.5% Triton X-100) for 6 days at room temperature (RT), and on the final day overnight (ON) at 4 ^◦^C. After staining, samples were washed with PBS and fixed again in 4% PFA for 24 hours at 4 ^◦^C. All samples were then washed again in PBS before being placed in 50% CUBIC-R+(M) (with dH20) (TCI, Cat#T3741) ON at RT and finally, placed in 100% CUBIC-R+(M) for a minimum of 3 days prior to imaging, to ensure samples were transparent.

Imaging was performed on a mesoSPIM light-sheet microscope [8] previously characterised [9] (Fig. 1a). Clarified left ventricular sections containing the injection site (Fig. 1b), were mounted in the inner cuvette with the epicardial surface oriented towards the detection objective (Fig. 1c,d).

**FIGURE 1.**
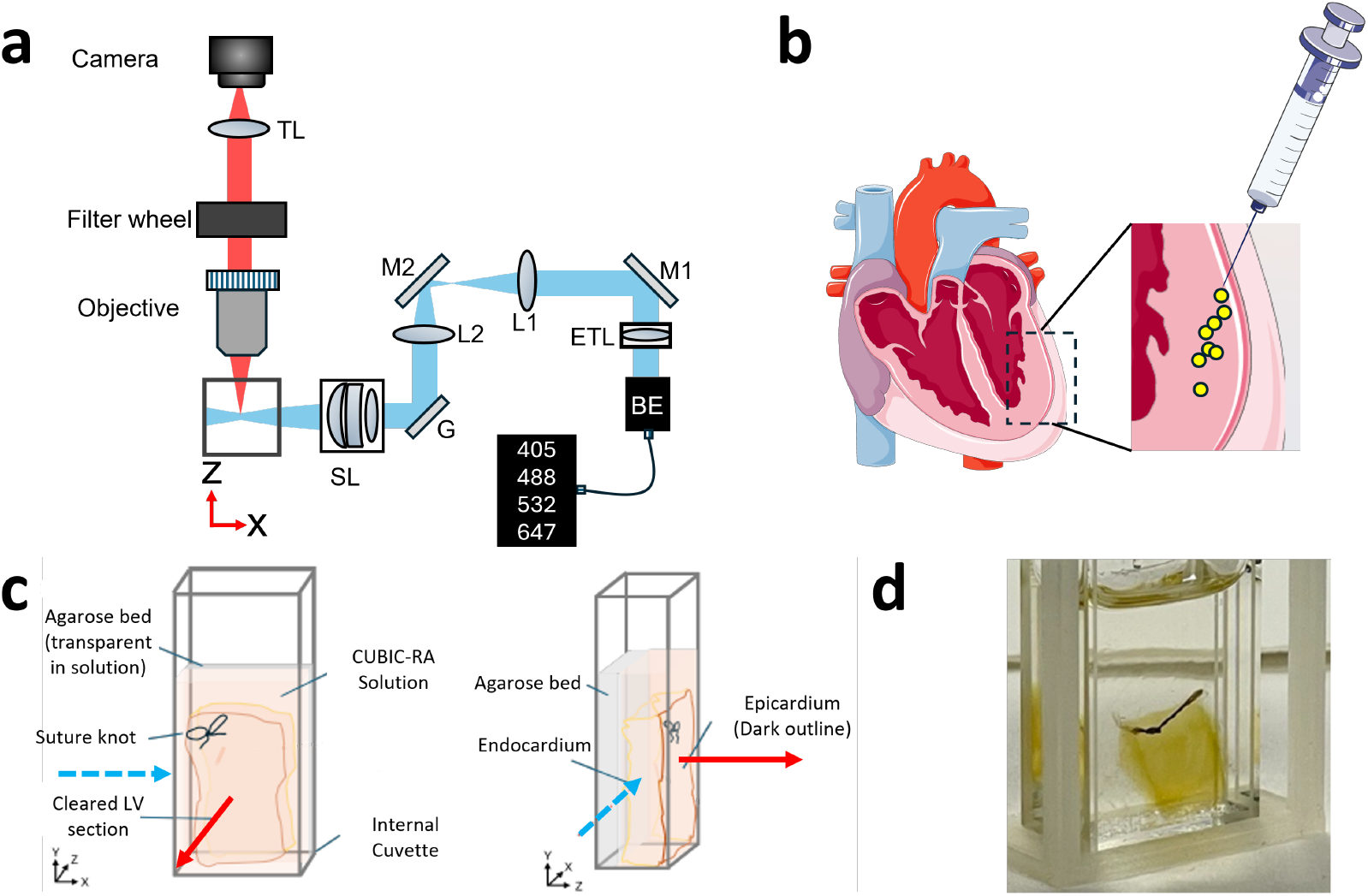
Imaging and sample preparation. **a)** Schematic of mesoSPIM light-sheet microscope. **b)** Illustration of spheroid injection into the left ventricle wall at a shallow angle relative to the epicardial surface. **c)** Cleared left ventricular section mounted in the inner cuvette for imaging. A suture marks the injection site. Blue dashed arrow indicates illumination axis and red solid arrow the detection axis. **d)** Photograph of a cleared left ventricular section mounted in the imaging cuvette with the injection site indicated by the suture.

### 2.2 Random Forest Classification and analysis

Pixel-wise classification of the raw image (Fig.2a) was performed in Ilastik [14] using a random-forest model comprising multiple decision trees. The features used to train the spheroid detecting classifier were raw intensity, Gaussian blur, Laplacian of Gaussian, Difference of Gaussians and Gaussian Gradient Magnitude filters (Fig.2b), capturing intensity, edge, and texture information. Each filter was applied with variances of *σ* = 1, 5, 10, 15, 20, 25, 30, and 50 to span the expected spheroid size range and smaller fragmented structures.

**FIGURE 2.**
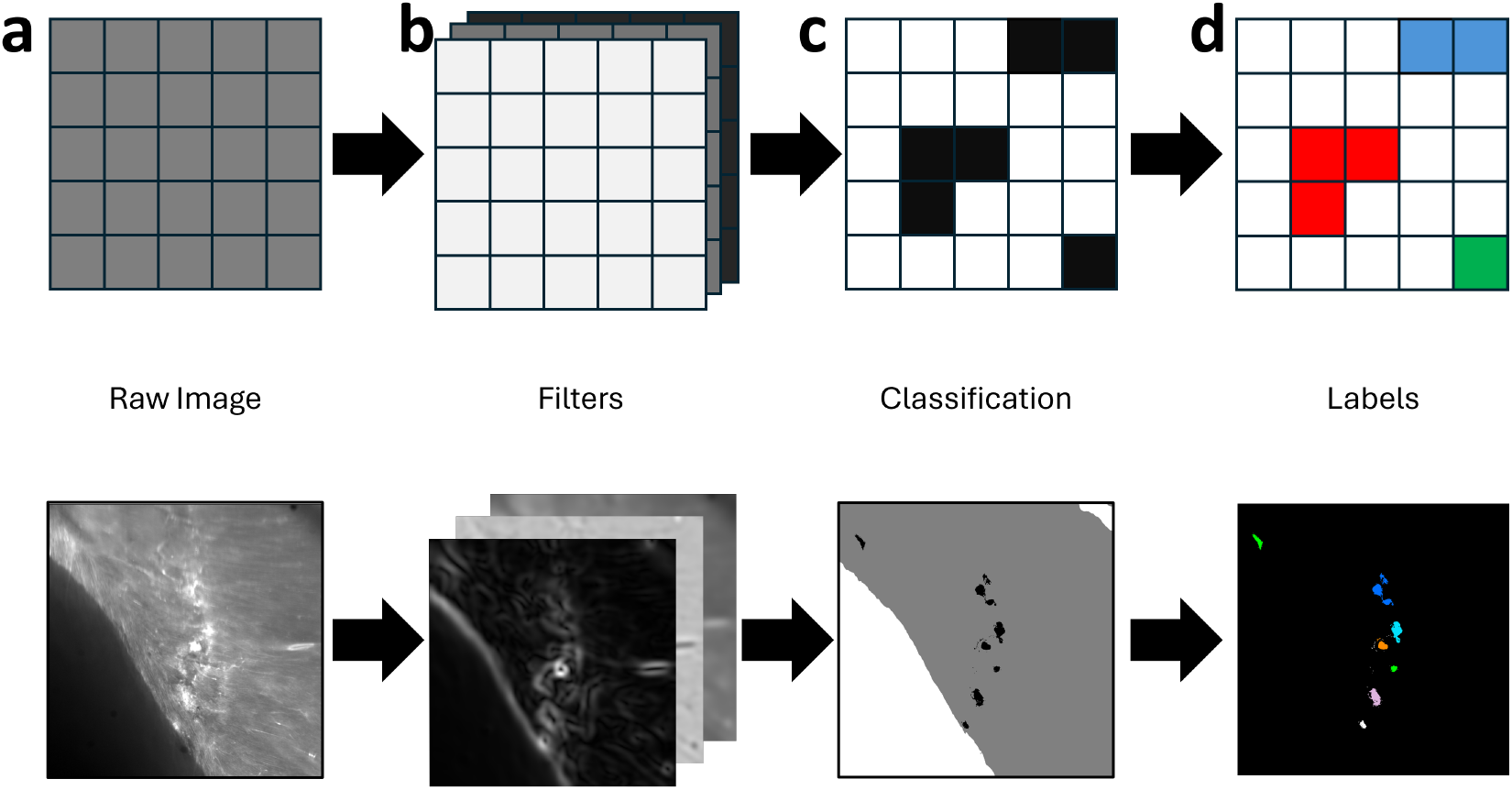
Segmentation pipeline shown schematically (top) and on representative experimental data (bottom). **a)** Raw input image. **b)** Feature extraction: multiscale filtered images used to train the classifier. **c)** Pixel-wise classification into spheroids, tissue and background classes. **d)** Object reconstruction: connected component grouping produces uniquely labelled spheroids.

Manual annotations were generated and reviewed by cardiovascular experts. Separate classifiers were trained for the hiPSC and H9C2 data sets using three classes: spheroid, surrounding tissue, and background. Each classifier was trained on a single representative WGA volume, selected for variation in structures and signal intensities. The training images were sparsely annotated to prevent overfitting. A similar number of annotations was made for each of the pixel classes to prevent overtraining on one class, resulting in a final classifier which would favour the labelling of pixels as the majority class and exhibit a poor accuracy in identifying the less represented class in the training data. Testing was performed on one hiPSC and two H9c2 WGA volumes. The classifier output is a three-class probability map assigning each pixel to spheroid, rabbit heart tissue or background (Fig.2c).

Probability maps were thresholded at 0.5 and converted to discrete labels. Connected-component analysis [16, 17, 18] (2-connectivity) grouped voxels into individual spheroid objects. This resulted in a dataset in which each spheroid was identified and uniquely labelled (Fig.2d). A size threshold on the number of voxels in an object allowed the filtering of the vast majority of false positives.

From this labelled dataset, several descriptive statistics parameters were extracted. The centroid, calculated by taking the mean of the positions of the constituent voxels, was taken to be the position of the spheroid. Spheroid volume was calculated by multiplying the number of voxels by voxel volume and the radius was then calculated assuming a spherical geometry. A three dimensional tree of spheroid positions was created using SciPy [16] to query their mutual distance [19], defined here as the distance from each spheroid to its nearest three neighbours. The epicardial surface was extracted from the tissue mask by marking the first voxel along the imaging axis. Spheroid depth was then computed as the Euclidean distance from each centroid to this surface, providing quantitative localisation of injected spheroids relative to the epicardium.

All analysis was done on a HP Zbook Firefly G9 Mobile Workstation with a 12th Gen Intel(R) Core(TM) i7-1280P processor (1.80 GHz) and 32gb of RAM.

## 3 RESULTS AND DISCUSSION

Using meso-scale light-sheet microscopy, we imaged the spheroid injection track beneath the epicardial surface of the rabbit left ventricle. Although identical labels were used for injected spheroids and host myocardium, spheroids were readily distinguished by morphology and cell density in both the SYTOX (Fig. 3a) and WGA channels (Fig. 3b). A supervised classifier generated spheroid probability maps across both hiPSC and H9c2 datasets (Fig 3c). Generating a probability map took around an hour per image for a stack size of a few gigabytes. However, this did vary and did not scale linearly for different image sizes. Applying a 0.5 probability threshold and connected-component grouping produced discrete three-dimensional spheroid objects distinct from surrounding tissue and background (Fig 3d). Feature extraction based on intensity, edge, and texture across Gaussian scales (*σ* = 1–50) enabled robust detection despite structural heterogeneity.

**FIGURE 3.**
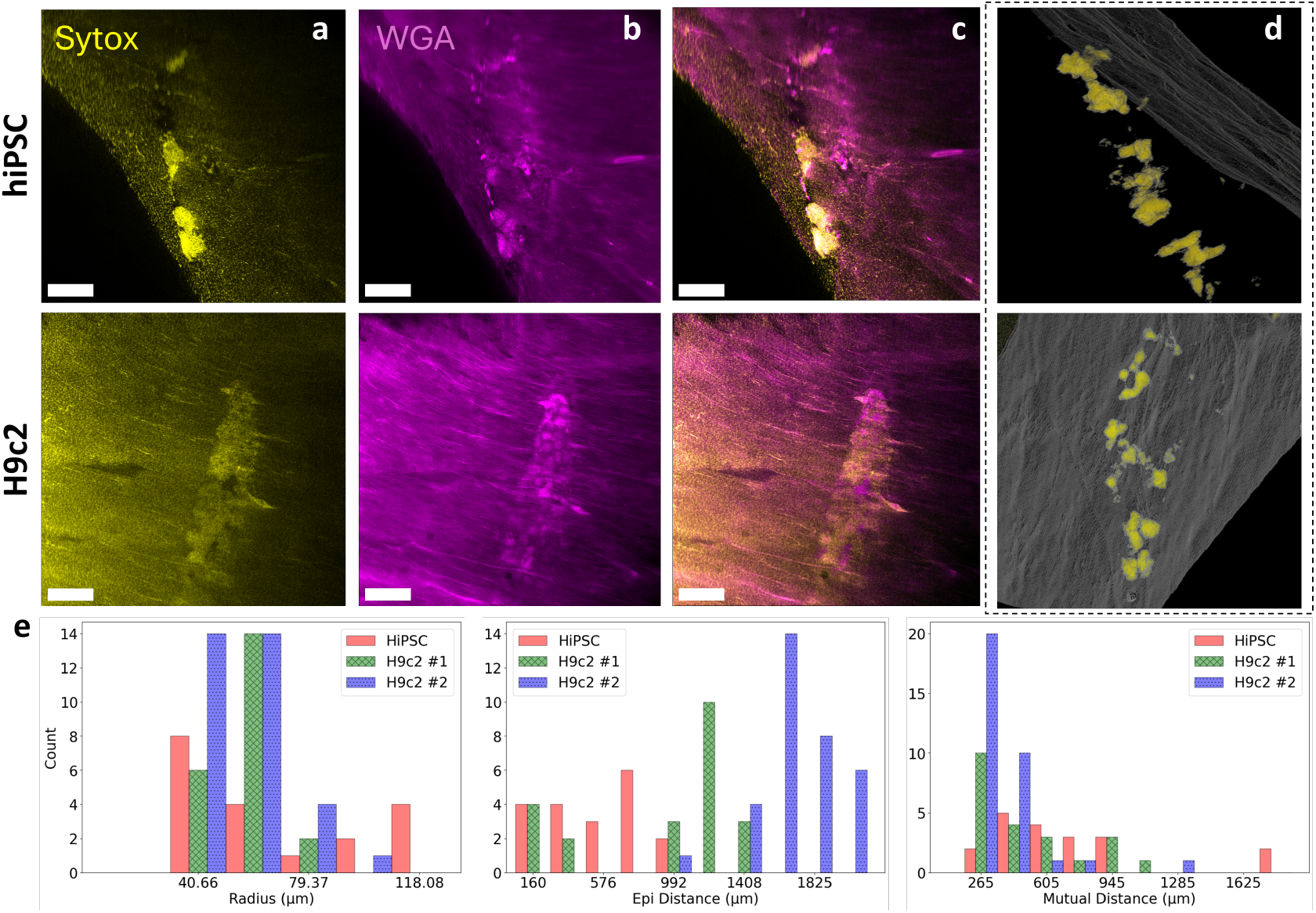
Volumetric imaging and quantitative analysis of injected spheroids. Scale bars = 600µm **a)** Representative Sytox channel frame from an hiPSC (top) and H9c2 (bottom) datasets. **b)** Corresponding WGA channel frame for hiPSC (top) and H9c2 (bottom) datasets. **c)** Dual channel overlays (Sytox/WGA) from hiPSC (top) and H9c2 (bottom) datasets. **d)** 3D rendering of segmented spheroids (yellow) relative to epicardial surface (grey) in the hiPSC sample. **e)** Histograms of extracted spatial metrics (radius, epicardial depth and nearest neighbour distance) across the three test volumes. Plots made using Matplotlib [20] and Seaborn [21]

Qualitative inspection confirmed consistent identification of spheroid cores and boundaries across imaging depths and signal levels. Three-class training (spheroid, surrounding tissue, background) reduced interface misclassification and prevented out-of-specimen labelling. Size filtering effectively removed small false positives while preserving intact spheroids. From randomly sampled frames which were also manually labelled, the calculated Dice scores between the classifier segmentation and the ground truth annotation were 0.761±0.130 (hiPSC classifier) and 0.731±0.117 (H9c2 classifier), indicating good agreement with manual annotation.

Object-level reconstruction enabled extraction of spheroid centroid, volume, and estimated radius for all detected spheroids ((Figure 3e)). A three-dimensional tree representation of centroids supported spatial analysis of inter-spheroid distribution. Segmentation of the epicardial surface from the tissue mask enabled computation of spheroid depth relative to the epicardium, providing quantitative localisation of injected clusters within the myocardium (Figure 3d).

### 3.1 Discussion

A key strength of the pipeline is the transition from voxel-level classification to object-level reconstruction and spatial quantification. By grouping classified voxels into connected components and extracting centroids, volumes, and estimated radii, the workflow enables automated derivation of structural metrics from whole-volume images. The implementation of a three-dimensional tree analysis for spheroid distribution permits the querying of spatial statistics, supporting quantitative assessment of spheroid spatial relationships. Additionally, extraction of the epicardial surface from the classified tissue mask provides a reproducible method for computing spheroid depth relative to the injection-facing surface, integrating anatomical context into the analysis.

The observed Dice scores (0.761 ± 0.130 for hiPSC and 0.731 ± 0.117 for H9c2) indicate good agreement between manual annotation and automated segmentation, particularly given the presence of numerous structures and variations in signal intensity and imaging depth. Qualitatively, it could be seen that the largest difference between the ground truth annotations and the segmentation was at the spheroid boundary, with the classifier tending to underestimate spheroid size by marking its edges as background tissue.

Although the Dice scores demonstrate good overlap overall, the high standard deviation between different frames (∼15% of absolute value) suggests that segmentation performance is influenced by local imaging conditions. Reduced performance in regions of low signal-to-noise or at irregular spheroid boundaries likely contributes to this spread. Nevertheless, the balanced annotation strategy and sparse labelling approach appear sufficient to prevent systematic bias toward any single class, supporting the generalisability of the trained classifiers within each dataset.

Compared with threshold-based segmentation approaches, the random forest framework accommodates non-linear relationships between features. However, it remains computationally lightweight and requires substantially less annotated data than deep learning–based alternatives, making it accessible for rapid deployment in a small laboratory setting.

Limitations include dependence on representative training volumes and manual annotation quality. Further work is required to assess cross-dataset transferability between samples which have been prepared differently. For example, the WGA channel tended to show spheroid-spheroid boundaries better, while the SYTOX channel showed the spheroid-tissue distinction better. Additionally, while Dice scores indicate good agreement, performance may decline in extreme low-contrast conditions or in cases of highly irregular spheroid morphology.

Overall, the presented approach provides a reproducible and scalable framework for automated three-dimensional spheroid segmentation and spatial analysis in cleared cardiac tissue, demonstrating that random forest pixel classification can effectively bridge detection and higher-order structural quantification in large volumetric imaging datasets.

## 4 CONCLUSION

We present a random forest–based pixel classification pipeline for automated three-dimensional segmentation and reconstruction of epicardially injected spheroids in optically cleared myocardial tissue. By integrating intensity, edge, and texture features across a broad range of spatial scales within Ilastik’s pixel classification framework, the method enables robust voxel-wise labelling and reliable object-level reconstruction from large light-sheet microscopy volumes.

The workflow combines sparse, expert-validated training annotations with probability thresholding, connected-component analysis, and size filtering to generate discrete spheroid objects suitable for volumetric and spatial quantification. Extraction of centroid position, volume, nearest-neighbour distances, and depth relative to the epicardium demonstrates the capacity of the approach to move from pixel classification to biologically relevant three-dimensional spatial metrics in an automated and reproducible manner.

Overall, this lightweight and accessible machine learning framework provides a scalable solution for spheroid detection in cleared cardiac tissue and establishes a pipeline for objective, high-throughput structural analysis of large volumetric imaging datasets.

## acknowledgements

Figure 1b provided by Servier Medical Art (https://smart.servier.com) licensed under CC BY 4.0 https://creativecommons.org/licenses/by/4.0/

